# Urine proteomic characterization of active and recovered COVID-19 patients

**DOI:** 10.1101/2023.03.12.532269

**Authors:** Jianping Sun, Jing Wei, Haibin Yu, Haidan Sun, Xiaoyan Liu, Yonghong Zhang, Chen Shao, Wei Sun, Jing Zhang, Youhe Gao

## Abstract

**Background:** The molecular changes in COVID-19 patients have been reported in many studies. However, there were limited attention has been given to the disease sequelae in the recovered COVID-19 patients.

**Methods:** Here, we profiled the urine proteome of a cohort of 29 COVID-19 patients in their disease onset and recovery period, including mild, severe, and fatal patients and survivors who recovered from mild or severe symptoms.

**Results:** The molecular changes in the COVID-19 onset period suggest that viral infections, immune response changes, multiple organ damage, cell injury, coagulation system changes and metabolic changes are associated with COVID-19 progression. The patients who recovered from COVID-19 still exhibited an innate immune response, coagulation system changes and central nervous system changes. We also proposed four potential biomarkers to monitor the whole progression period of COVID-19.

**Conclusions:** Our findings provide valuable knowledge about the potential molecular pathological changes and biomarkers that can be used to monitor the whole period of COVID-19.

## Background

The ongoing COVID-19 pandemic, caused by severe acute respiratory syndrome coronavirus 2 (SARS-CoV-2), has led to 754,367,807 confirmed cases and 6,825,461 deaths worldwide as reported by the World Health Organization (WHO) on 6, February 2023, with a tremendous impact on global health and economics. SARS-CoV-2 has been reported to infect multiple organs, including the lung, liver, heart, muscle, gastrointestinal tract, spleen, testis, thyroid and central nervous system[1-4]. Currently, most proteomic studies focus on the molecular changes in COVID-19 patients, such as the immune response, blood coagulation and fibrosis, as well as potential biomarkers for severity evaluation[4-8]. Only limited studies have revealed the molecular changes between active and recovered COVID-19 patients using serum proteomics[9,10]. As up to 87.4% of patients who have recovered from COVID-19 report persistent fatigue and dyspnea[11], there is still an urgent need for a comprehensive understanding of the molecular level changes throughout the entire course of COVID-19 patients.

Urine has been regarded as a promising biomarker source, as it is noninvasive, easily collected and can reflect multiple organ changes[12,13]. Notably, urine samples are less complex than blood samples and carry many proteins, peptides, and amino acids that have not been discovered in blood. Currently, a total of 6085 proteins have been reported to be identified in normal human urine[14]. The urine proteome has been applied to stratify patients with familial Parkinson’s disease[15] and to discriminate lung cancer patients from other common tumors[16]. In addition, the urine proteome has also been used to reflect the immune response in moderate and severe COVID-19 patients[17,18] and to identify potential biomarkers to predict patients with mild to severe symptoms[19]. However, the study of biomarkers to monitor the whole process of COVID-19 is still unavailable.

In this study, we presented a urinary proteomic analysis including the entire course of COVID-19 from the onset to the progression and recovery periods. This whole course included a total of six phases, namely, the mild phase (M), the severe phase (S), the fatal phase (F), the mild remission phase (MRE), the severe remission phase (SRE), and the severe return visit phase (SRT). The mass spectrometry-based, data-independent acquisition (DIA) quantitative proteomic approach was performed to analyze urine samples from healthy donors, COVID-19 patients and their recovery period. The altered urine proteins were classified into six categories: virus infection, immune response, organ damage, cell injury, coagulation system changes, and metabolic changes. We also analyzed our urine proteomic data in conjunction with the clinical serum indicators of the COVID-19 patients. Finally, we propose candidate monitoring urine biomarkers for the entire course of COVID-19. These biomarkers can be used not only as diagnostic biomarkers of COVID-19 but also as potential prognostic biomarkers to predict disease recovery. Our data provide valuable knowledge of the molecular changes in the whole period of COVID-19, including the recovery phase, and shed light on the potential biomarkers for monitoring the entire disease course.

## Methods

### Experimental Design and Statistical Rationale

COVID-19 patients were recruited in Beijing Youan Hospital from January to February 2020. All work performed in this study was approved by the Beijing Youan Hospital Ethics Committee (LL-2020-030-K). Written informed consent was obtained from the patients. Patients were diagnosed with COVID-19 based on the Chinese Government Diagnosis and Treatment Guideline (Trial 5th version) published by the National Health Commission of China (National Health Commission, 2020)[20]. The healthy donors were recruited from health care workers at Beijing Youan Hospital, and none of them had previously been infected with SARS-CoV-2.

SARS-CoV-2-positive patients were enrolled in this study after diagnosis. According to the abovementioned diagnosis standards, these recruited COVID-19 patients were classified into three subgroups: 1) mild-mild symptoms mainly manifested with symptoms of fever, nonpneumonia or mild pneumonia; 2) severe-fulfilled any of the following three criteria: respiratory distress, respiratory rate (R ≥ 30 times/min); mean oxygen saturation (≤ 93%, resting state); arterial blood oxygen partial pressure/oxygen concentration (PaO2/FiO2 ≤ 300 mmHg); 3) fatal - fulfilled any of the three criteria: respiratory failure and required mechanical ventilation; shock incidence; admission to the ICU.

In this study, we collected urine samples from a cohort of 29 COVID-19 patients, including 12 patients diagnosed with mild (M) symptoms, 16 patients diagnosed with severe (S) symptoms and 2 patients with fatal (F) outcomes. The 12 mild patients included a total of 19 urine samples for proteomic analysis, namely, 6 mild onset period samples (M), 11 mild remission samples (MRE) and 2 mild return visit samples (MRT). The 16 severe patients included a total of 31 urine samples for proteomic analysis, namely, 10 severe onset period samples (S), 16 severe remission samples (SRE) and 5 severe return visit (SRT) samples. The two fatal patients included 3 fatal onset period urine samples (F). The remission urine samples were taken after the patients tested negative for SARS-CoV-2. Midstream first morning urine was collected from these subjects. After collection, the urine was immediately centrifuged at 5,000 g for 30 min at 4 °C to remove the cell debris and then sterilized at 56 °C to inactivate the virus.

### Proteomics sample preparation

Urine samples were prepared using the FASP method [21]with some modifications. Briefly, 4 ml urine from each specimen was reduced with 20 mmol/L dithiothreitol (DTT, Sigma) at 95 °C for 5 min and then alkylated by 50 mmol/L iodoacetamide (IAA, Sigma) for 45 min in the dark. Then, the urine samples were precipitated with six volumes of ethanol at -20 °C overnight. The pellets were dissolved in lysis buffer (8 mol/L urea, 2 mol/L thiourea 50 mmol/L Tris, and 25 mmol/L DTT). The protein amount of each sample was measured using the Bradford method.

A total of 100 μg of protein from each sample was loaded onto a 10 kDa filter device (Pall, Port Washington, NY, USA). After being washed two times with urea buffer (UA, 8 mol/L urea, 0.1 mol/L Tris-HCl, pH 8.5) and 25 mmol/L NH_4_HCO_3_ solutions, these samples were digested with trypsin at 37 °C for 14 h. The digested peptides were desalted using Oasis HLB cartridges (Waters, Milford, MA, USA) and dried by vacuum evaporation (Thermo Fisher Scientific, Bremen, Germany). The peptide concentration was measured using the BCA method.

### Spin column peptide fractionation

Digested peptides were dissolved in 0.1% formic acid and diluted to 0.5 μg/μL. A pooled sample (96 μg, 1.5 µg of each sample) from 64 samples was loaded onto a high-pH, reversed-phase fractionation spin column (84868, Thermo Fisher Scientific) to generate a spectral library. Briefly, the spin column was equilibrated with acetonitrile and 0.1% formic acid solution. Then, the sample solution was loaded on the column to elute the “flow-through” fraction, while the water was loaded on the column to elute the “wash” fraction. Next, a step gradient of 8 increasing acetonitrile concentrations (5, 7.5, 10, 12.5, 15, 17.5, 20 and 50% acetonitrile) in a volatile high-pH elution solution was added to the column to elute these peptides into eight different gradient fractions. Finally, the fractionated peptides were evaporated and resuspended in 20 μl of 0.1% formic acid. Two microliters of each fraction was loaded for LC-DDA-MS/MS analysis.

### LC–MS/MS analysis

LC–MS/MS data acquisition was carried out on a Orbitrap Fusion Lumos mass spectrometer coupled with an Easy-nLC 1200 system (both from Thermo Scientific). The iRT reagent (Biognosys, Switzerland) was added at a concentration of 1:20 v/v in all peptide samples to calibrate the retention time of extracted peptide peaks. For analysis, 1 μg of peptide from each sample was loaded into a C18 trap column (75 μm * 2 cm, 3 μm, C18, 100 Å) and then separated with a reversed-phase analytical column (100 μm * 100 cm, 2 μm) at a flow rate 1 μL/min with a 120 min gradient (buffer B: 2%–6% for 1 min, 6%–10% for 23 min, 10%–20% for 67 min, 20%–28% for 7 min, 28%–95% for 20 min, and 95%–5% for 2 min). The LC settings were the same in both DDA-MS and DIA-MS modes to maintain a stable retention time. Ten fractions from spin column separation were analyzed with mass spectrometry in DDA mode to generate the spectral library. The MS data were acquired in high-sensitivity mode. A full MS scan was acquired in a 350-1,500 m/z range with the resolution set to 120,000. The MS/MS scan was acquired in Orbitrap with a resolution of 30,000. The HCD collision energy was set to 32%. The AGC target was set to 5e4, and the maximum injection time was 45 ms.

Then, 54 individual urine samples were analyzed in DIA-MS mode. The variable isolation window of the DIA method with 26 windows was developed for DIA acquisition. Positive ion mode was set to 3000 V. The full scan was set at a resolution of 120,000 with a m/z range from 350 to 1,200, and the DIA scan was set at a resolution of 30,000. The maximum injection time was 50 ms. The HCD collision energy was set to 32%. To ensure the quality of the data, the pooled peptides of all samples were used to ensure the stability of the instrument. A single pooled DIA analysis was run among after every 7-9 samples to serve as the technical replicate quality control. Different groups of samples were acquired in a random order to reduce system bias during mass spectrometry analysis.

### Mass spectrometry data processing

The MS data of the fractionated pools (DDA MS data, 10 COVID-19 pool urine and 15 human normal pool urine) and the single-shot subject samples (DIA MS data, 64 COVID-19 samples and 8 quality control samples) were used to generate a DDA library and direct DIA library, respectively, which were computationally merged into three cohort-specific hybrid libraries using Spectronaut (v14.0.200409.43655, Biognosys AG). The searched database included the human SwissProt database (released in May 2019, containing 20,421 sequences), the SARS-CoV-2 proteome database containing 14 protein sequences and the 11 peptide iRT sequence. Searches used carbamidomethylation as a fixed modification and acetylation of the protein N-terminus and oxidation of methionines as variable modifications. The trypsin/P proteolytic cleavage rule was used, permitting a maximum of 2 missed cleavages and a minimum peptide length of 7 amino acids. The Q-value cutoffs for both library generation and DIA analyses at the peptide and protein levels were set to 0.01.

A total of 8 QC samples and 64 DIA samples were imported into Spectronaut Pulsar X with default settings. The optimal XIC extraction window was determined according to the iRT calibration strategy. Cross-run normalization was performed to calibrate systematic variance in LC–MS performance, and local normalization based on local regression was used [22]. Protein inference was performed using the implemented IDPicker algorithm [23]. All results were then filtered by a Q value less than 0.01 (corresponding to an FDR of 1%). The peptide intensity was calculated by summing the peak areas of the respective fragment ions for MS2. Protein intensity was calculated by summing the respective peptide intensity.

### Statistical analysis

To analyze the proteomic data, we first used the sequential-KNN method to impute the missing value of QC samples[24]. Missing values larger than 50% were removed for later analysis. The QC samples were used as technical replicates to evaluate the stability of mass spectrometry. Then, the 64 DIA samples were imputed as the healthy control group, the disease group, and the recovery group.

For later analysis, we normalized the protein abundance by log2 transformation for P value analysis. Differential urinary proteins in the six periods of COVID-19 were compared with the healthy control group using the following criteria: fold change ≥2 or ≤0.5; comparison between two groups were conducted using a two-sided, unpaired t test; and P values of group differences were adjusted by the Benjamini and Hochberg method[25]. Group differences resulting in adjusted P values <0.05 were considered statistically significant. All results are expressed as the mean ± standard deviation.

### Bioinformatics analysis

Protein interaction network analysis was performed using the STRING database (https://string-db.org/cgi/input.pl) and visualized by Cytoscape (V.3.7.1) [26]. The canonical pathways as well as disease and function categories were enriched using IPA software (Ingenuity Systems, Mountain View, CA). The bubble figures were visualized by the Wu Kong platform (https://www.omicsolution.org/wkomics/main/).

### Biomarker screening

The potential urine biomarkers for monitoring were selected according to the following criteria: i) differential proteins were present in all three active phases, in the M and S phases, or in the S and F phases of COVID-19; ii) differential proteins were upregulated/downregulated gradually in the active phase but changed reversely in the recovery period; iii) differential proteins were not significantly different in the recovery stages compared with the healthy control; and iv) the potential monitoring biomarkers were associated with the molecular changes of COVID-19 patients.

## Results

### Study design and clinical characteristics

We collected urine samples from a cohort of 29 COVID-19 patients, including 12 patients diagnosed with mild (M) symptoms, 16 patients diagnosed with severe (S) symptoms and 2 patients with fatal (F) outcomes. Healthy control urine samples were collected from 11 healthy (H) donors at Beijing Youan Hospital. The patients in the M and S groups survived COVID-19 and were discharged from the hospital. Of note, the 12 mild patients included a total of 19 urine samples for proteomic analysis, namely, 6 mild onset period samples (M), 11 mild remission samples (MRE) and 2 mild return visit samples (MRT). The 16 severe patients included a total of 31 urine samples for proteomic analysis, namely, 10 severe onset period samples (S), 16 severe remission samples (SRE) and 5 severe return visit (SRT) samples. The two fatal patients included 3 fatal onset period urine samples (F) (Figure 1). The remission urine samples were taken after the patients tested negative for SARS-CoV-2.

**Figure 1.**
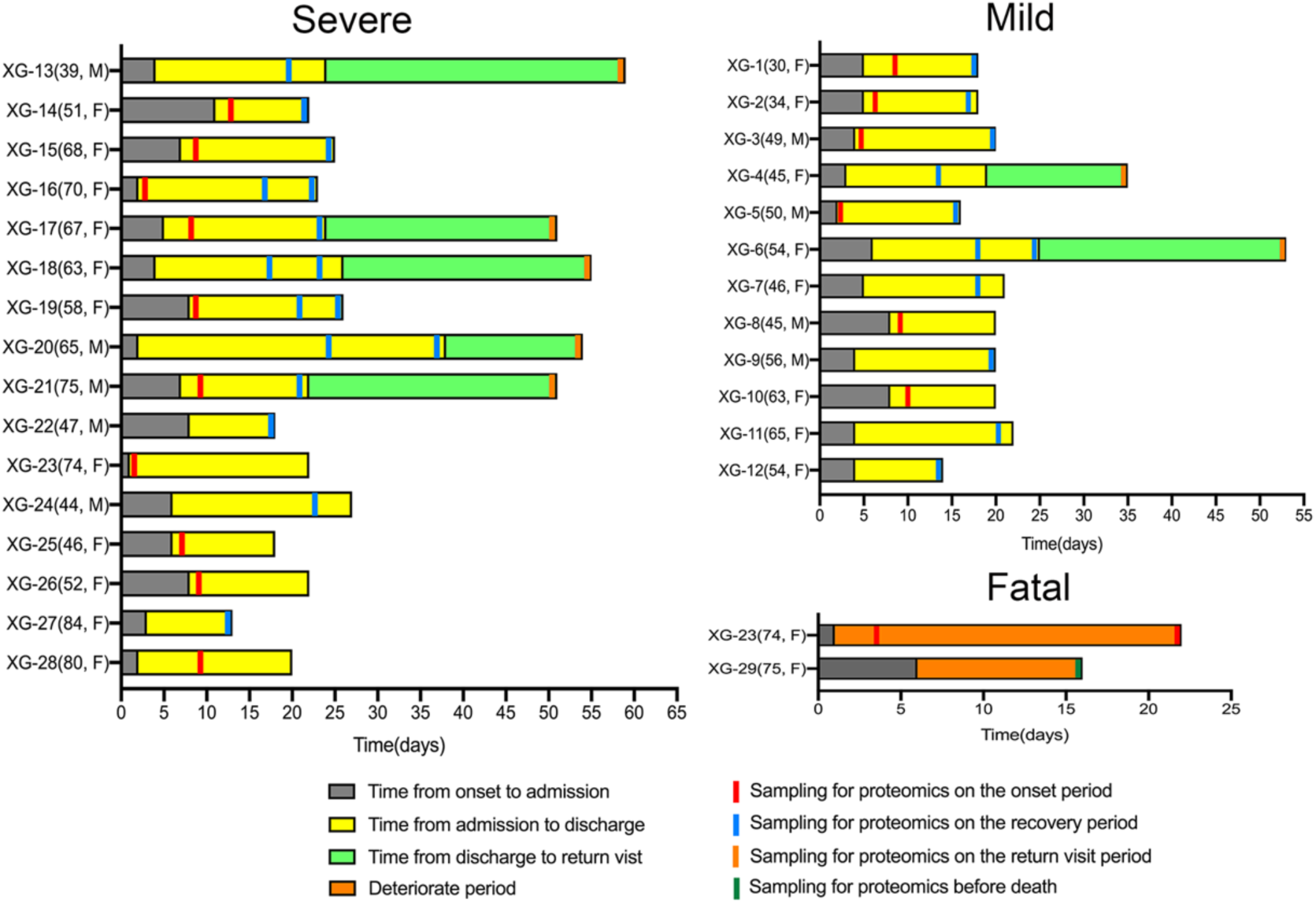
Timeline of the disease course and sample acquisition of COVID-19 patients, including 12 mild patients, 16 severe patients and two fatal patients. Further details are presented in Table 1 and Table S1.

The detailed patient descriptions in their onset and recovery phases are shown in Table 1. In addition, we also analyzed 28 clinical measurements of the COVID-19 patients in their onset and recovery period (Figure S1, Table S1). These clinical indicators cover liver function, kidney function, immune and inflammatory changes, myocardial function and coagulation functional changes. As shown in Figure S1, we noticed that the clinical indicators of liver function, such as AST, ALT, DBIL and TBIL, were upregulated in the active period but downregulated in the recovery period. Immune and inflammatory indicators, such as N and WBC, were upregulated in the active period but downregulated in the recovery period. LYMPH exhibited the reverse change compared with N and WBC. Coagulation function indicators such as TT and myocardial function indicators such as CK-MB and TnI were upregulated in the active phase of COVID-19 but downregulated in the recovery phase. When compared with the M or S patients, the fatal patients showed significant suppression of lymphocyte count, as well as increases in neutrophil count, WBC, AST, CK-MB, MYO, TnI, TT and HGB. When compared with the COVID-19 patients in their onset period, we noticed that the PLT, LYMPH, MONO and AST showed the reversed change in their recovery phase.

**Table 1.**
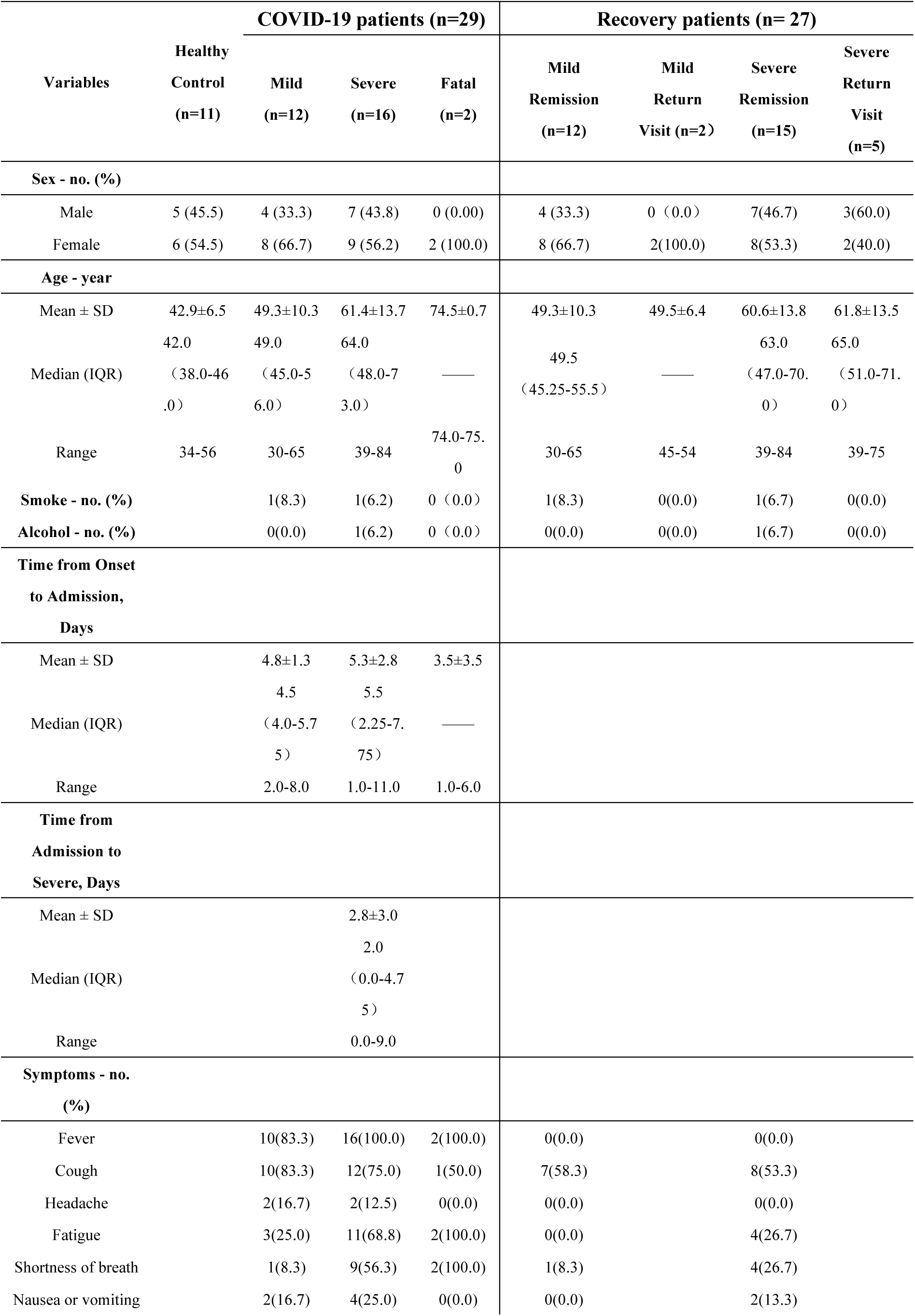
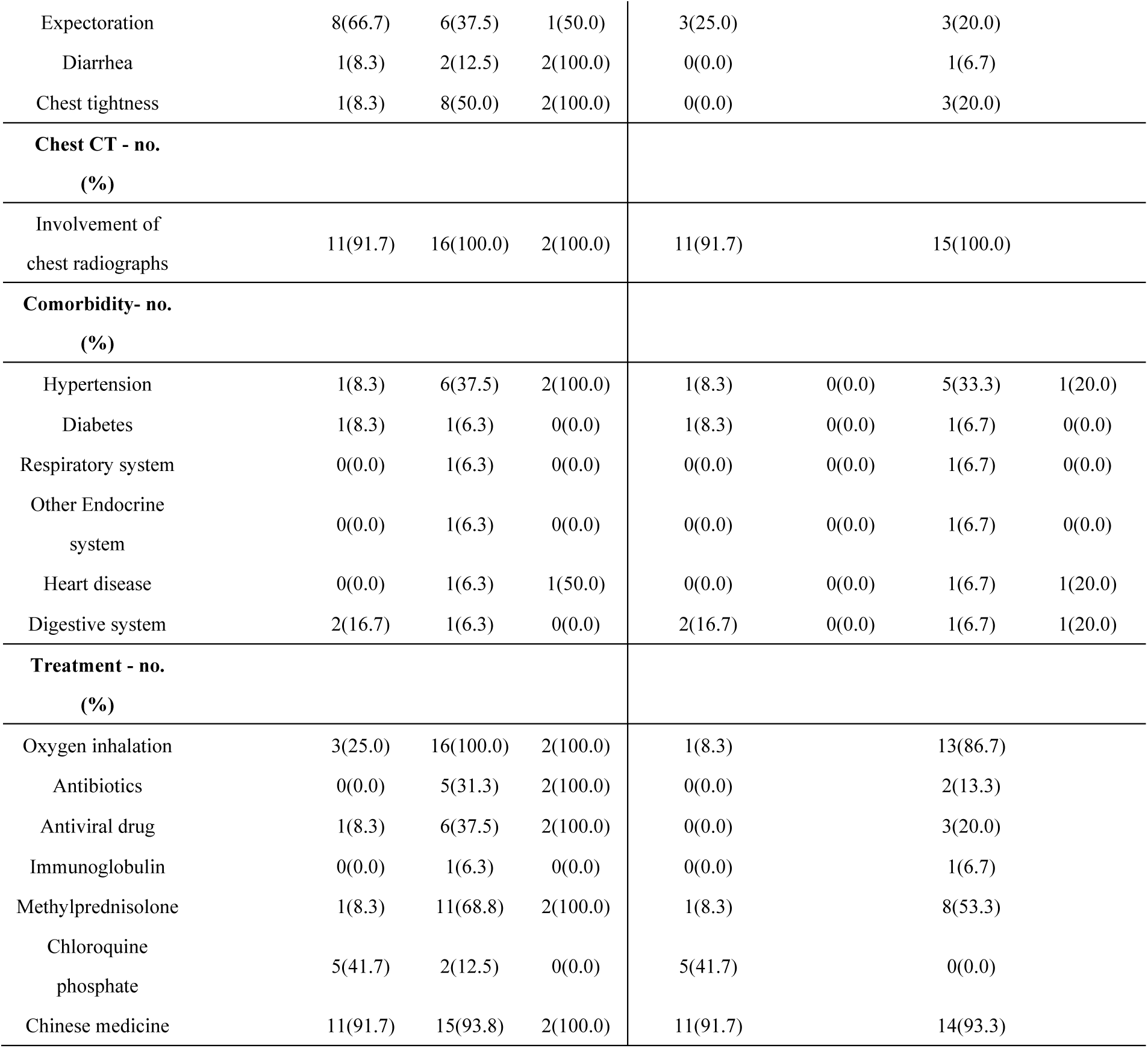
Demographics and baseline characteristics of COVID-19 patients in their active and recovery periods.

### Quantitative urine proteome profiling of COVID-19 patients in the disease and recovery stages

We used a data-independent acquisition (DIA) strategy and liquid chromatography/tandem mass spectrometry (LC–MS/MS) to analyze the urine samples. A total of 3367 protein groups were identified from 64 urine samples with a false discovery date (FDR) of less than 1% at both the peptide and protein levels. A total of 2950 protein groups were identified with at least two peptides. The number of protein groups in each sample is presented in Figure S2A. No SARS-CoV-2 proteins were identified from 64 urine samples. The median coefficient of variance (CV) and the correlation of QC samples indicated the great consistency and reproducibility of our data (Figure S2B, S2C). A total of 2574 protein groups of the healthy group, COVID-19 group and recovery group were retained, as their missing values were less than 50% (Table S2). The t-distributed stochastic neighbor embedding (t-SNE) analysis of the 2574 protein groups showed clear stratification between the COVID-19 patients, the recovery group and the healthy group, while the recovery group was more similar to the healthy control (Figure 2A).

**Figure 2.**
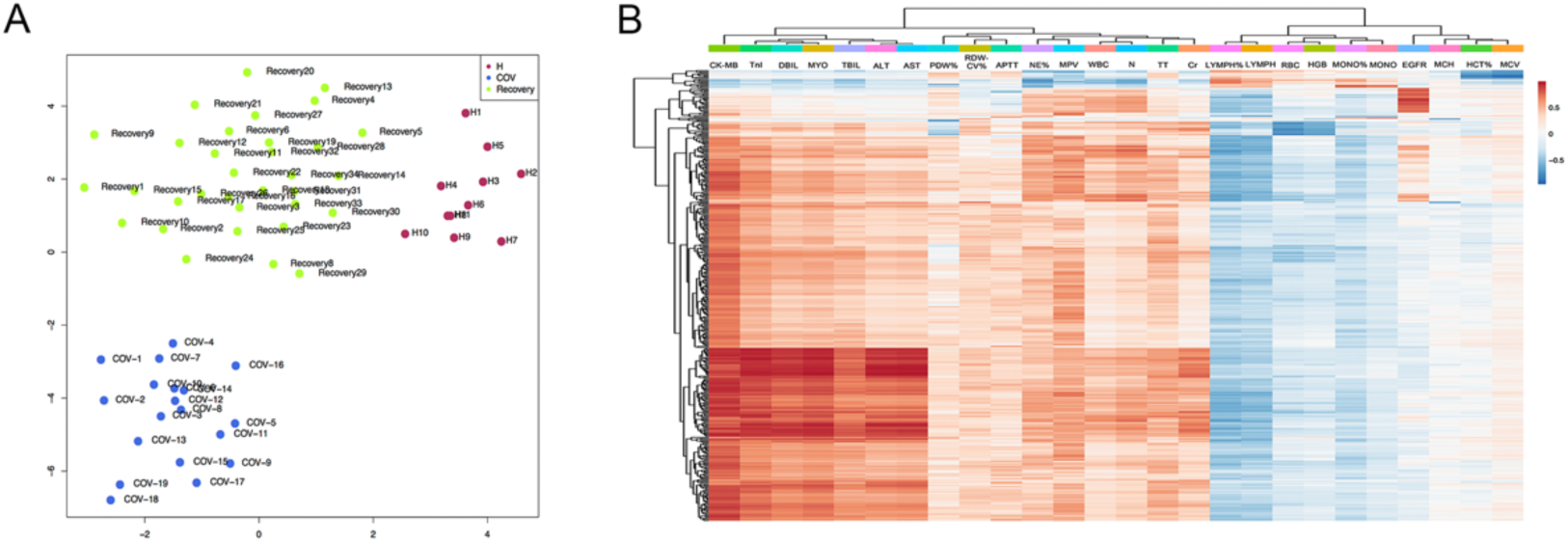
Quality control of proteomic data and the correlation with clinical indicators. (A) t-distributed stochastic neighbor embedding (t-SNE) of urine samples using 2574 measured urine protein groups. (B). Pearson correlation analysis between the differentially expressed proteins in the onset period and 28 clinical indicators.

Differential proteins were all compared with the healthy group with fold change≥2 and Benjamini–Hochberg (B-H) adjusted p value<0.05. Finally, there were 291, 601, 987, 245, 336 and 213 differentially expressed proteins in the M, S, F, MRE, SRE and SRT groups, respectively (Table S2). Pearson correlation analysis between the differentially expressed proteins in the M, S and F phases and the 28 clinical indicators was performed. Finally, 456 differentially expressed proteins showed a significant correlation with 26 clinical indicators (Figure 2B). These clinical indicators, including the liver, kidney, heart, immune response and coagulation changes, were subgrouped into two clusters. Some clinical indicators, including myocardial function, liver function, kidney functions and a few immune functions, exhibited a positive correlation with our differential urinary proteins, while some other clinical indicators, such as LYMPH, MONO, RBC and HGB, showed a negative correlation with our differential urinary proteins. Our results indicated that our urine proteome changes are highly associated with clinical indicators, such as abnormal liver and kidney function, myocardial abnormalities, disturbances in blood coagulation and abnormal immune or inflammatory changes.

The differential urinary proteins of the six groups were then enriched by the IPA database to reveal the molecular landscape associated with SARS-CoV-2 infections. The enriched canonical pathways as well as the disease and biofunctions were classified into six clusters to reveal the whole process changes in COVID-19 patients in their onset and recovery period. These six clusters included viral infection changes, immune response changes, multiple organ damage, cell injury, coagulation system changes and metabolic changes.

### Viral infection changes

The IPA database was used to enrich the related viral infection changes of six periods of COVID-19. The viral infection molecular changes in COVID-19 patients in their onset and recovery periods are presented in Figure 3A. The majority of these molecular changes, such as viral infection, infection of cells, infection by SARS/RNA/coronavirus and the coronavirus replication pathway, are active in the disease phase of COVID-19. Pathways such as viral infection, infection by SARS coronavirus/RNA virus/coronavirus and the coronavirus replication pathway were active in the three active phases of COVID-19, and the active extent was associated with the severity of COVID-19.

**Figure 3.**
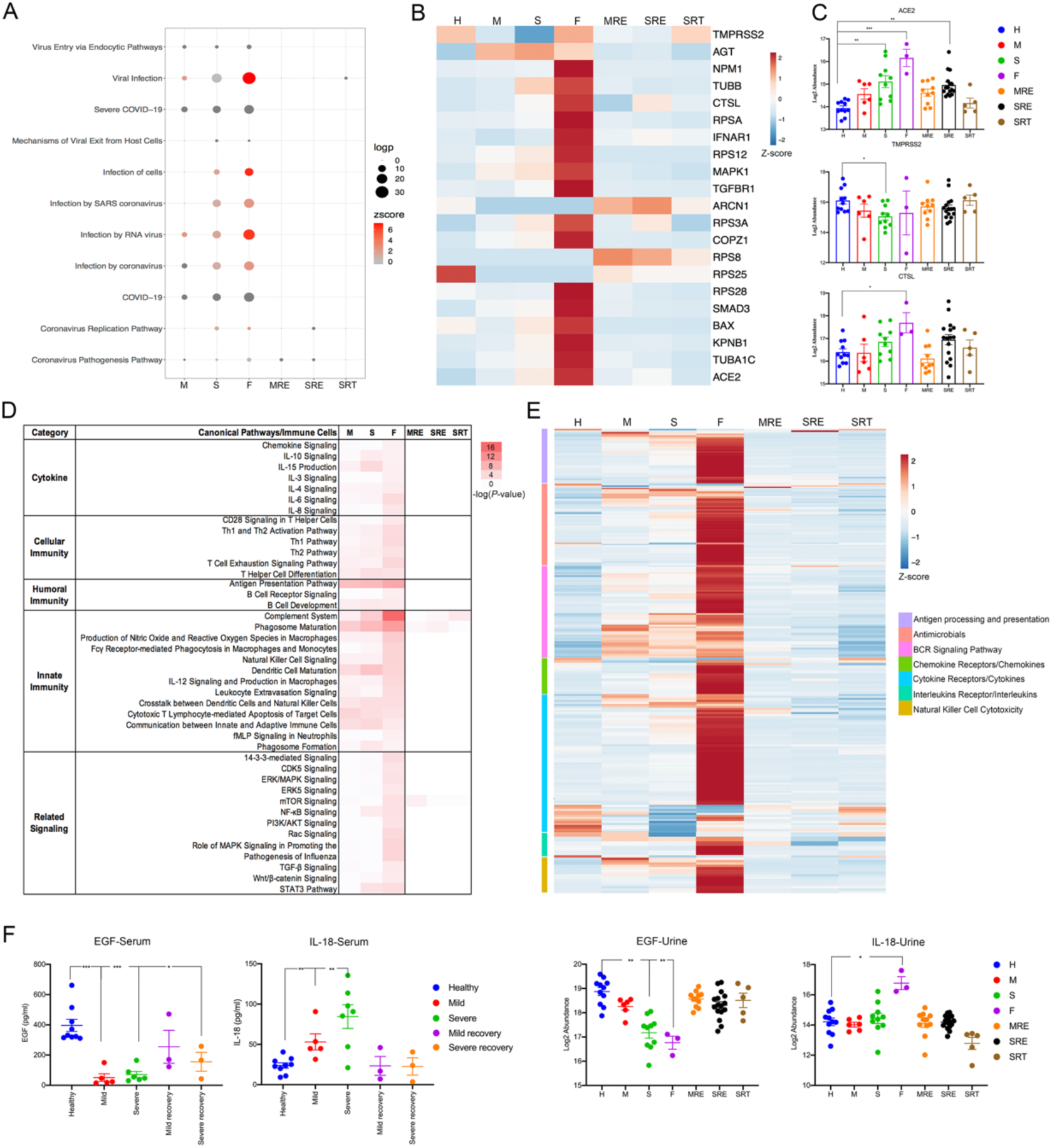
Viral infection and immune response changes in COVID-19 patients in the onset and recovery periods. (A) The dysregulated pathways and disease/biofunctions associated with viral infections. Pathway and disease/biofunction enrichment analyses were performed using differential proteins in the six phases (M, S, F, MRE, SRE, and SRT) using IPA. The size of the circle represents the -log10(p value), and the color represents the Z score by IPA. (B) Heatmap of 21 differentially expressed proteins associated with the coronavirus replication pathway and the coronavirus pathogenesis pathway. The average abundance of proteins in the six phases of COVID-19 was calculated and visualized as a heatmap. (C) Protein expression of ACE2, TMPRSS2 and CTSL in the six phases (M, S, F, MRE, SRE, and SRT) of COVID-19 and healthy controls. The y-axis represents the protein expression transformed by the log2 abundance. Pairwise comparison of each protein between the six phases and the healthy control was performed with Student’s t test. The cutoff of the significant phase was set at a B-H adjusted p value<0.05 and FC>2 or <0.5. B-H adjusted p value: *p<0.05; **p<0.01; ***p<0.001. (D) The dysregulated pathways associated with immune response changes. Pathway enrichment analysis was performed using differential proteins in the six phases (M, S, F, MRE, SRE, and SRT) using IPA. The color represents the -log10(p value) by IPA. (E). Heatmap of the immune response classified into seven clusters using the ImmPort database. The average abundance of proteins in the six phases of COVID-19 was calculated and visualized as a heatmap. (F). Clinical parameters and the protein expression levels of EGF and IL-18 in the active and recovery periods of COVID-19 patients. The y-axis represents the protein expression transformed by the log2 abundance or the concentration in serum. The significance indicated by the asterisks. For the clinical parameters, the unpaired two-sided Welch’s t test was performed (p value: *, < 0.05; **, < 0.01; ***, < 0.001). For urine protein expression levels, significance was set at B-H adjusted p value<0.05 and FC>2 or <0.5. B-H adjusted p value: *p<0.05; **p<0.01; ***p<0.001.

There were 21 differential proteins involved in the coronavirus replication pathway and the coronavirus pathogenesis pathway (Figure 3B). We noticed that cathepsin L1 (CTSL), which is involved in the coronavirus replication pathway, was significantly upregulated in the fatal phase of COVID-19. Transmembrane protease serine 2 (TMPRSS2) and angiotensin-converting enzyme 2 (ACE2) were enriched in coronavirus replication and the coronavirus pathogenesis pathway (Figure 3C). Of note, various 40S ribosomal proteins such as RPSA, RPS12, RPS3A, RPS8, RPS25 and RPS28, were enriched in the coronavirus pathogenesis pathway, indicating that these 40S ribosomal proteins may contribute to the inhibition of the translation system in COVID-19 patients.

### Immune response changes

We found that most immune cells, such as phagosomes, macrophages, monocytes, NK cells, T cells, leukocytes, B cells and neutrophils, were enriched in the disease period of COVID-19, while phagosome maturation was still present in the SRE phase of patients. In addition, immune responses, such as cytokines, cell and humoral immunity, innate immunity and some immune-related signaling pathways, were found to be associated with the progression of COVID-19, while innate immunity, such as the complement system, was still present in the SRT phase of severe patients when they recovered from COVID-19 (Figure 3D). In addition, we classified the immune-related urine differential proteins into seven clusters according to the ImmPort database (Figure 3E). Most of these immune-associated differential urine proteins were upregulated gradually, especially in the fatal phase of COVID-19, and decreased in the recovery stages. These results indicated that the immune response is active in the onset period of COVID-19 patients, which is consistent with disease progression and may still be present when they test negative for SARS-CoV-2 infections.

As the cytokine storm contributes to the majority of deaths from COVID-19, we then focused on the common pathways associated with cytokines. A total of 15 common differential proteins associated with the cytokine pathways were identified both in the IPA and ImmPort databases (Figure S3A). We also evaluated a total of 48 cytokine concentrations in the serum of these COVID-19 patients during both the disease period and the recovery period. Serum samples from fatal patients were lacking, as most of the serum samples were collected from COVID-19 patients in the early period of the pandemic outbreak, and the most important job was to save patients’ lives at that time. As shown in Figure S3B, the levels of the representative eotaxin, IFN-α2, interleukins (IL-1α, IL-5, IL-6, IL-10), monocyte chemoattractant protein-1 (MCP-1) and monocyte chemoattractant protein-3 (MCP-3) increased significantly in the M and/or S phases of COVID-19 and were downregulated in the recovery periods.

### Changes in multiple organs

To estimate the potential organ damage of COVID-19, we enriched differential proteins associated with multiple organs, including brain, heart, lung, liver, kidney and muscle damage, according to the IPA database (Table S3I).

We first compared the differential urine proteins associated with damage to the heart, kidney, lung and liver to the corresponding autopsy results of COVID-19 patients [4] and selected urine proteins with the same fold change (Figure 4A). We also noticed that the neurological system showed a strong association with the disease period of COVID-19 patients and may still be present in their recovery period (Figure 4B). Finally, digestive system and muscle damage were also found according to our results. Digestive system damage appeared not only in severe and fatal patients with COVID-19 but also in the recovery SRE phase. Taking these potential damages into consideration, we found that most of them appeared in the severe and fatal phases, while some of them were still present in the SRE phase, indicating that potential organ damage may still be present in the recovery phase of COVID-19.

**Figure 4.**
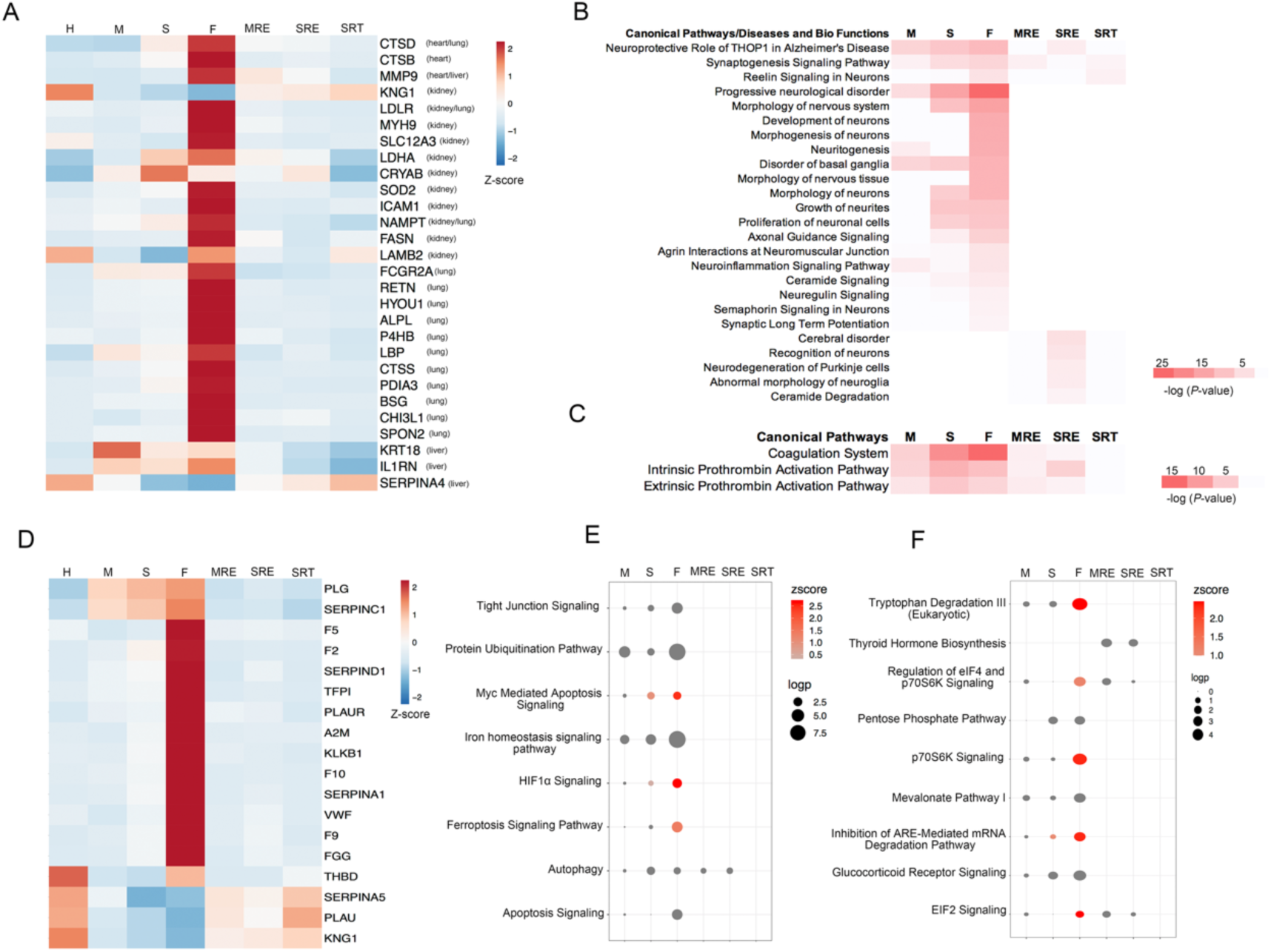
Multiple organ damage, cell injury, coagulation system changes and metabolic changes in COVID-19 patients in the onset and recovery periods. (A) Heatmap of common differential urine proteins identified in the autopsy results of the heart, kidney, lung and liver. The average abundance of proteins in the six phases of COVID-19 was calculated and visualized as a heatmap. (B) The dysregulated pathways and disease/biofunctions associated with neurological system changes. Pathway and disease/biofunction enrichment analyses were performed using differential proteins in the six phases (M, S, F, MRE, SRE, and SRT) using IPA. The color represents the -log10(p value) by IPA. (C) The dysregulated pathways associated with coagulation system changes. Pathway enrichment analysis was performed using differential proteins in the six phases (M, S, F, MRE, SRE, and SRT) using IPA. The color represents the -log10(p value) by IPA. (D). Heatmap of 18 differentially expressed proteins associated with the coagulation system. The average abundance of proteins in the six phases of COVID-19 was calculated and visualized as a heatmap. (D) The dysregulated pathways associated with cell injury. Pathway enrichment analysis was performed using differential proteins in the six phases (M, S, F, MRE, SRE, and SRT) using IPA. The size of the circle represents the -log10(p value), and the color represents the Z score by IPA. (E) The dysregulated pathways associated with metabolic changes. Pathway enrichment analysis was performed using differential proteins in the six phases (M, S, F, MRE, SRE, and SRT) using IPA. The size of the circle represents the -log10(p value), and the color represents the Z score by IPA.

### Coagulation system changes cell injury

Coagulation system changes appear not only in the disease period of COVID-19 patients but also in their recovery phase (Figure 4C). Various enriched functional lists, such as the coagulation system, thrombus, blood clot and degranulation of blood platelets, were associated with the progression of COVID-19. Notably, we noticed that the coagulation system, intrinsic/extrinsic prothrombin activation pathway and accumulation of blood cells were still present in the SRE phase of patients, indicating that abnormal coagulation may still exist after a negative nucleic acid test. There were 18 differential proteins participated in coagulation system changes (Figure 4D). Interestingly, four differential proteins, urokinase-type plasminogen activator (PLAU), kininogen-1 (KNG1), plasma serine protease inhibitor (SERPINA5) and thrombomodulin (THBD), were downregulated in the disease period but upregulated in the recovery period.

### Cell injury and Metabolic changes

The majority of pathways associated with cell injury, such as tight junction signaling, the protein ubiquitination pathway, Myc-mediated apoptosis signaling and the ferroptosis signaling pathway, showed a strong association with the severity of COVID-19 (Figure 4E). HIF-1α signaling was active in the severe and fatal phases of patients in our results.

Changes in amino acids, nuclear acids, glucose and fatty acids were observed in the disease period of patients. We found that amino acid degradation appeared in most of the amino acid metabolism pathways. Of note, thyroid hormone biosynthesis appeared only in the MRE and SRE phases in COVID-19 patients, while the mevalonate pathway I was found to exist in fatal COVID-19 patients. Lipid metabolism plays a crucial role in viral infections, while viruses can use the host lipid machinery to support their life cycle and to impair the host immune response. Therefore, the altered expression of mevalonate pathway-related genes induced by SARS-CoV-2 will ensure their survival and spread in host tissue[27] (Figure 4F).

### Potential urine biomarkers to monitor the whole period of COVID-19

To identify potential specific urine proteins monitoring the whole period of COVID-19, we clustered the 1186 differential proteins in the disease active period into 9 significant discrete clusters with the quantified normalized abundance through mFuzz[28]. According to the previous criteria, we chose Cluster 5 and Cluster 7 for further selection (Figure 5A). Finally, superoxide dismutase [Cu-Zn] (SOD1) and complement factor B (CFB) in Cluster 7 and kininogen-1 (KNG1) and pro-epidermal growth factor (EGF) in Cluster 5 were selected as potential biomarkers for monitoring the whole process of COVID-19 (Figure 5B). SOD1 participates in the superoxide radical degradation pathway, which is categorized as a metabolic change, while CFB participates in the complement system, which is categorized as an immune response in COVID-19 patients. KNG1 and EGF were associated with coagulation changes and immune response changes, as we mentioned before. We also identified another 24 differential proteins, as they also exhibited great differentiation ability in the three active stages of COVID-19 but without systematic molecular functions, as we presented above (Figure S4).

**Figure 5.**
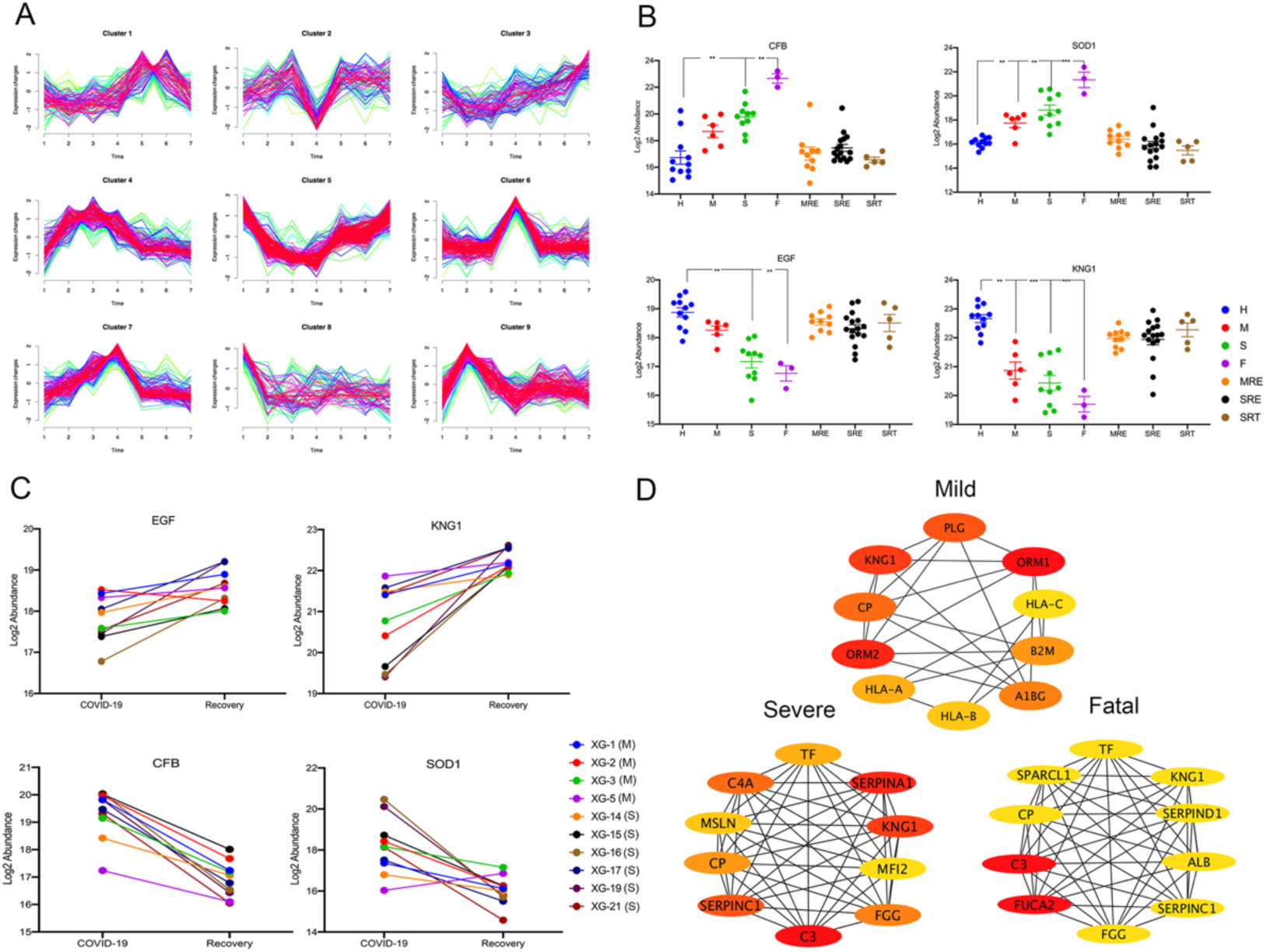
Potential monitoring biomarkers in the COVID-19 active and recovery periods. (A) Identification of specific clusters in COVID-19 patients. A total of 1181 differentially expressed proteins in the COVID-19 active period were clustered using mFuzz into significant discrete clusters to illustrate the relative expression changes of the 1186 differentially expressed proteins. The groups in proteomics data: 1: Healthy; 2: Mild; 3: Severe; 4: Fatal; 5: MRE; 6: SRE; 7: SRT. (B) Protein expression of CFB, SOD1, EGF and KNG1 in the six phases (M, S, F, MRE, SRE, and SRT) of COVID-19 and healthy controls. The y-axis represents the protein expression transformed by the log2 abundance. Pairwise comparison of each protein between the six phases and the healthy control was performed with Student’s t test. The cutoff of the significant phase was set at a B-H adjusted p value<0.05 and FC>2 or <0.5. B-H adjusted p value: *p<0.05; **p<0.01; ***p<0.001. (C) Validation of CFB, SOD1, EGF and KNG1 expression in 10 COVID-19 patients, including 4 mild and 6 severe patients. The y-axis represents the protein expression transformed by the log2 abundance. (D) The ten hub proteins of differential proteins in the mild, severe and fatal phases using the MCC method in the cytoHubba module of CytoScape.

The four candidate monitoring biomarkers (SOD1, CFB, EGF, and KNG1) were then evaluated in 4 mild and 6 severe patients, as these 10 patients had a complete timeline in their onset and recovery period (Figure 5C). The expression of KNG1 and EGF was upregulated, while SOD1 and CFB expression was downregulated in the recovery phase of the 4 mild and 6 severe patients compared with their onset period, indicating great potential monitoring functions as biomarkers.

Finally, we also tried to identify potential diagnostic urine biomarkers in the M, S and F phases of COVID-19 patients. The STRING database was used to generate the PPI network of the DEPs in these three disease phases. Then, the MCC method in the cytoHubba module of CytoScape was used to identify the top 10 hub proteins in the three disease phases (Figure 5D). The area under the curve (AUC) was calculated according to the ROC curve. Interestingly, we found that KNG1 appeared as a hub protein in all three disease periods, with an AUC larger than 0.98 (Figure S5). This result indicated that KNG1 can not only be a potential diagnostic urine biomarker in the M phase of COVID-19 but can also be used as a disease monitoring biomarker in the whole period of COVID-19.

## Discussion

The urine proteome has been applied in various aspects of COVID-19, such as to uncover the immune response[29] and identify signaling pathways and therapeutic options[30]. In this study, we focused on the urine proteome changes of the entire course in COVID-19 patients from their onset to the progression and recovery periods, and try to identify potential biomarkers to diagnose and predict COVID-19 recovery.

The urine proteome can reflect viral infection changes. We noticed that cathepsin L1 (CTSL), which is involved in the coronavirus replication pathway, was significantly upregulated in the fatal phase of COVID-19. CTSL is considered the serine protease of SARS-CoV-2 in the endosomal pathway and mediates the cleavage of the S1 subunit of the coronavirus surface spike glycoprotein[31]. Since this cleavage is necessary for coronavirus entry into human host cells, virus and host cell endosome membrane fusion, and viral RNA release for the next round of replication, the inhibitor of CTSL has been proven to be an effective treatment option for COVID-19 by blocking virus entry[31-33]. CTSL has also been reported to be upregulated in the lung as a potential receptor for virus entry[4]. Transmembrane protease serine 2 (TMPRSS2) and angiotensin-converting enzyme 2 (ACE2) were enriched in coronavirus replication and the coronavirus pathogenesis pathway. It has been reported that SARS-CoV-2 uses the SARS-CoV receptor ACE2 for entry and the serine protease TMPRSS2 for S protein priming, while the TMPRSS2 inhibitor might constitute a treatment option for clinical use by blocking virus entry[34,35]. In addition, as SARS-CoV-2 nonstructural protein 1 (Nsp1) was reported to inhibit protein translation of model human and SARS-CoV-2 messenger RNAs (mRNAs) by specifically binding to the small (40S) ribosomal subunit[36-38], we therefore suppose that these 40S ribosomal proteins may contribute to the inhibition of the translation system in COVID-19 patients.Interestingly, we still noticed that the coronavirus replication pathway, the coronavirus pathogenesis pathway and viral infection were enriched in the three recovery phases of COVID-19 patients despite the negative SARS-CoV-2 test. We therefore suppose that this may be due to the extremely limited number of viruses that cannot reach the standard of the nucleic acid test. It is worth noting that there were still some patients who were reported to test positive when they recovered from COVID-19.

The urine proteome can reflect immune response changes. The Interleukin-1 receptor antagonist protein (IL1RN) was reported to be a potential therapeutic method in acute hyperinflammatory respiratory failure in COVID-19 patients [39]. Similarly, the placenta growth factor (PGF) was reported to have mitogenic, angiogenic, and immunoregulatory properties, enhancing cellular survival, and is involved in host defensive mechanisms, which play an important role in adapting the host to newer infections, making it a promising therapeutic candidate against the COVID-19 virus[40]. Tumor necrosis factor receptor superfamily member 1A (TNFRSF1A) is an immune-based biomarker associated with mortality in COVID-19 patients[41]. Interleukin-18 (IL-18) was reported to be associated with other inflammatory markers and to reflect disease severity[42]. The antagonist of interleukin-6 receptor (IL-6R) can effectively block the IL-6 signal transduction pathway, making it an effective drug for patients with severe COVID-19[43-45]. Azurocidin (AZU1) was found to be significantly altered in naso-oropharyngeal samples of SARS-CoV-2 patients[46]. IL-1α is a ubiquitous and pivotal proinflammatory cytokine that is central to the pathogenesis of numerous conditions characterized by organ or tissue inflammation, including the complex multifactorial COVID-19[47]. IL-6 and IL-10 have been reported as predictors for the fast diagnosis of patients with a higher risk of disease deterioration[48]. MCP-1 and MCP-3 were reported to be biomarkers associated with the severity of COVID-19, which can be related to the progression and risk of death in COVID-19 patients[49,50]. Notably, we found that four urine differential proteins were common to these 48 cytokine-associated proteins, namely, IL1RN, IL-18, EGF and fractalkine (CX3CL1). IL-18 overproduction was reported to participate in the cytokine storm of COVID-19, causing an exaggerated inflammatory burden and leading to tissue injury[51]. Interestingly, we found that EGF was downregulated in the disease period of COVID-19 both in serum and urine samples but upregulated in the recovery phase. Epidermal growth factor receptor (EGFR) was reported to be upregulated significantly in lung fibrosis tissue[52], while inhibiting EGFR signaling may prevent an excessive fibrotic response to SARS-CoV disease[53]. In addition, EGFR was also regarded as one of the pivotal targets of puerarin in the treatment of COVID-19[54] (Figure 3F). We therefore suppose that the downregulation of EGF in serum and urine may be associated with the severe progression of COVID-19.

We also found that urine proteome may reflect potential organ damage. Vasogenic edema with mild infiltration of inflammatory cells and myocardial cell edema were the major pathological changes that were observed in the hearts of COVID-19 patients. As an inflammatory mediator, matrix metalloproteinase-9 (MMP9) was found to be elevated in the heart and liver of COVID-19 patients and was reported to facilitate matrix degradation and contribute to inflammatory cell invasion[55]. In addition, MMP-9 was also reported as an early indicator of respiratory failure in COVID-19 patients[56]. Lung fibrosis is the most significant pathologic injury in the lungs of COVID-19 patients, accompanied by fibrous exudation, hyaline membrane formation and alveolar space collapse[57]. The immunoglobulin gamma Fc receptor II-a (FCGR2A) was reported to be associated with COVID-19 susceptibility and upregulated in the lung tissue of COVID-19 patients[4,58].The enriched disease and biofunctions associated with brain damage, such as amyloidosis, progressive encephalopathy, Alzheimer’s disease (AD), dementia and brain lesions, were increasingly significant, consistent with the progression of COVID-19. The SARS-CoV-2 virus was reported to accelerate the pathogenesis of prion disease[59]. In addition, it has been reported that AD, Parkinson’s disease (PD), and dementias are associated with more complicated clinical courses and poorer outcomes in COVID-19 patients[60]. The above results indicated that COVID-19 may deteriorate some brain diseases, such as AD and dementia. Notably, there is increasing evidence that SARS-CoV-2 contributes to a number of neurological issues, including anosmia, seizures, stroke, confusion, encephalopathy, and total paralysis[61,62]. In addition, COVID-19 patients in the intensive care unit (ICU) have neurological deficits at a higher risk of mortality[63], and when they leave the ICU, they may potentially be at higher risk for long-term residual neuropsychiatric and neurocognitive conditions[64,65]. Additionally, the enriched axonal guidance signaling in our results has been reported to promote the rapid spread of the virus from neurons to neurons[66]. As ACE2 was found to be highly expressed in gastrointestinal epithelial cells, gastrointestinal symptoms such as diarrhea, vomiting, and abdominal pain were also reported in some COVID-19 patients[67]. Muscle damage, such as the cell movement of muscle cells, motor dysfunction or movement disorder, and neuromuscular disease, were found to be associated with COVID-19 progression in our results. It has been reported that the COVID-19 pandemic has the potential to disproportionately and severely affect patients with neuromuscular disorders[68]. In addition, myalgia was reported to be described among the common symptoms of COVID-19 after fever, cough, and sore throat, and the duration of myalgia may be related to the severity of COVID-19[69].

The urine proteome can reflect cell injury. Autophagy, the protein ubiquitination pathway and apoptosis signaling are common mechanisms of cell death. Targeting the mechanisms of autophagy was reported as an alternative therapeutic approach for COVID-19, while the clinical decision to inhibit or activate autophagy should depend on the underlying context of the pathological timeline of COVID-19[70]. The ubiquitin modifications can regulate the innate immune response by affecting regulatory proteins, either by altering their stability via the ubiquitin proteasome pathway or by directly regulating their activity. As the intrinsic deubiquitinating activities of papain-like protease (PLP), which is encoded by coronaviruses, antagonize the host’s immune response, PLP inhibitors were reported to show promising results as an antiviral therapy in vitro[71]. Furthermore, it has been reported that SARS-CoV-2 infection not only activates apoptosis in lymphocytes, causing lymphopenia[72], but also activates caspase-8 to trigger cell apoptosis and inflammatory cytokine processing in lung epithelial cells[73]. Notably, the ferroptosis signaling pathway was found to be upregulated in fatal COVID-19 patients and did not appear in the recovery phase of COVID-19 patients, which was consistent with the blood single-cell results of COVID-19 patients[74]. Tight junction signaling was associated with the severity of COVID-19 in our results. As tight junctions were reported to exist between epithelial cells in many organs, such as the lung, intestine, kidney and brain[75-77], the disruption of tight junction complexes is the major cause of epithelial barrier breakdown during virus infection[78]. It has been reported that the activation of the HIF-1α signaling pathway under severe hypoxic conditions will induce the activation of proinflammatory cytokine expression and the subsequent inflammation process and cytokine storm phase of COVID-19, which is consistent with our results[79].

The urine proteome can reflect coagulation system changes. PLAU was reported as a thrombolytic agent to positively resolve existing thrombi by accelerating the activity of plasmin[80]. According to the UniProt database, kininogens are inhibitors of thiol proteases, and high-molecular-weight (HMW)-kininogen inhibits the thrombin- and plasmin-induced aggregation of thrombocytes. We therefore suppose that it is reasonable for the downregulation of KNG1 in the disease period of COVID-19 patients. As a member of the serine protease inhibitor family (serpin), antithrombin was reported to be an important protease inhibitor of thrombin and factor Xa[81]. Additionally, THBD was reported to play a vital role in maintaining intravascular patency due to its anticoagulant, anti-inflammatory, and cytoprotective properties[82].

The urine proteome can reflect metabolic changes. The inhibition of the ARE-mediated mRNA degradation pathway, regulation of eIF4 and p70S6K signaling, EIF2 signaling and p70S6K signaling were reported to be translation-associated pathways[4].The reduction in tryptophan degradation was reported to be significantly associated with respiratory severity and correlated with inflammation in COVID-19 patients[83].Thyroid hormones play a key role in modulating metabolism and the immune system. Although thyroid function tests in survivors of COVID-19 have been reported to return to normal at baseline[84], it cannot be ignored that thyroid function still needs to be considered according to our results. The dysregulated glucose metabolism in severe COVID-19 suggests that SARS-CoV-2 pathogenesis involves a novel interplay with glucose metabolism[85]. Glucose metabolism was found to be dysregulated in the disease period of COVID-19 patients in our results. It has been reported that the upregulation of glucocorticoid receptor alpha appeared in COVID-19 patients[86]. In addition, in a SARS-CoV-2 infection asymptomatic ferret model, the pentose phosphate pathway was also observed in nasal wash samples using a metabonomic method[87].

There are some limitations in our study. First, all of the samples were collected from the patients in the early period of COVID-19 outbreak. Due to the priority to save patients’ lives at that time, the number of urine samples were limited and small. A larger number of samples were needed to validate these potential biomarkers in future research. Another possible drawback of this study is that different therapeutic strategies used during the treatment of different patients might affect the results, although the protein alterations uncovered here are quite consistent in different cohorts. Finally, the detailed roles of biomarker proteins such as EGF, KNG1, CFB and SOD1 require further investigation or experimentally validated.

## Conclusion

In summary, our study provides the whole system molecular changes of six aspects in COVID-19 patients ranging from the mild period to the severe return visit period (Figure 6). The coronavirus replication/pathogenesis pathway, the immune response changes and the coagulation system changes are associate with the severity of COVID-19 disease. The brain damage as well as the central nervous system changes are also enriched based on our urine proteome data. We also provide potential urine protein biomarkers for monitoring the whole process of COVID-19 including the different stages of active and recovery period. Our finding may provide useful monitoring and therapeutic strategies in the ongoing battle against COVID-19 pandemic.

**Figure 6.**
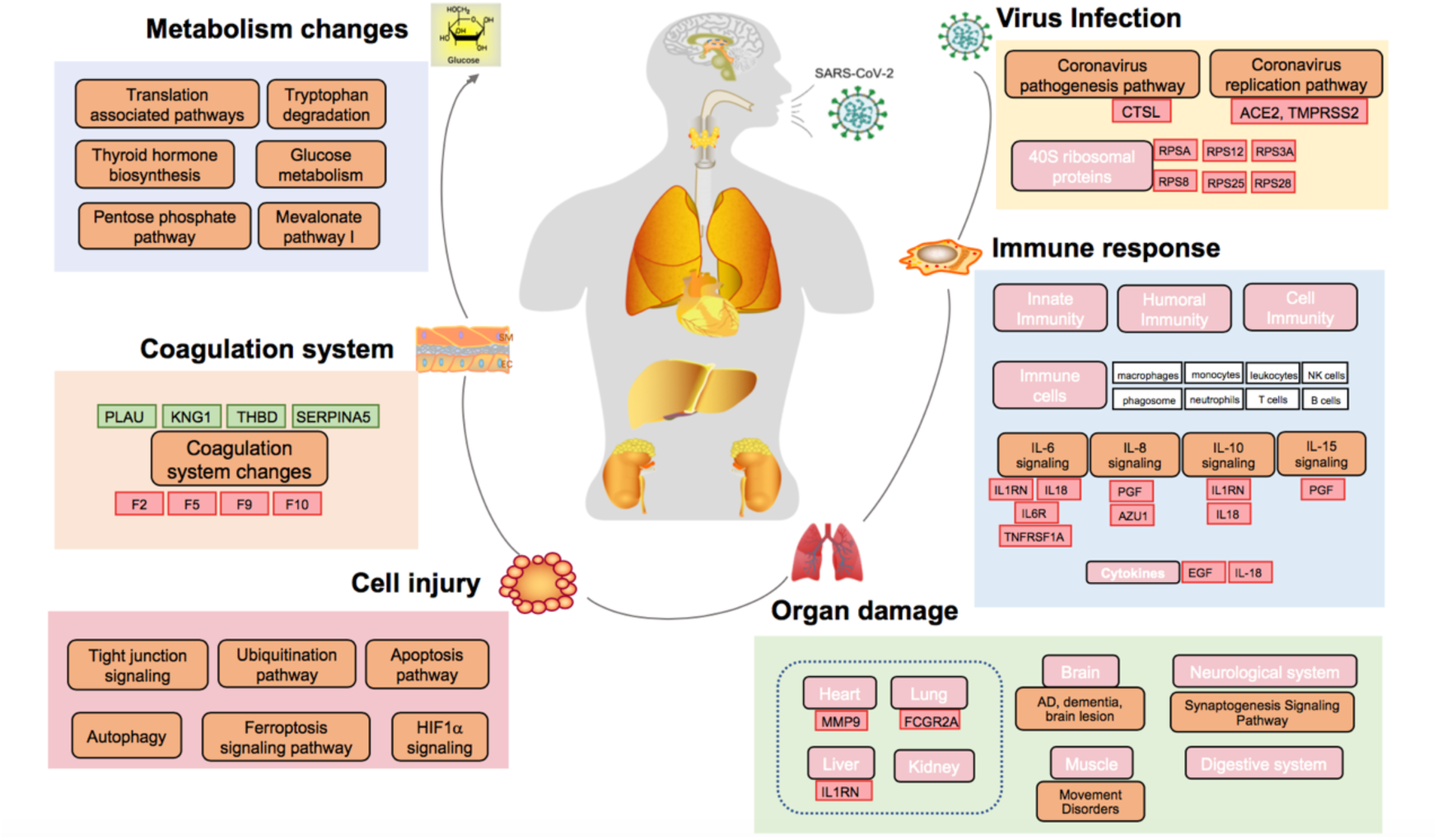
Dysregulated proteins and pathways in six aspects of functional changes in COVID-19. A hypothetical system view of multiple functional changes in response to SARS-CoV-2 infection. Red boxes: upregulated proteins; green boxes: downregulated proteins; orange/pink box: molecular processes; white box: related immune cells.

## List of abbreviations

LC-MS/MS: liquid chromatography-tandem mass spectrometry;
BMI: body mass index;
CI: confidence interval;
CV: coefficient of variation;
eGFR: estimated glomerular filtration rate;
FABP5: fatty acid binding protein 5;
FASP: filter-aided sample preparation;
QC: quality control;
RI: reference interval;
ROC: receiver operation curve;
COVID-19: Corona Virus Disease 2019;
SARS-CoV-2: severe acute respiratory syndrome coronavirus 2;
MRE: mild remission;
SRE: severe remission;
SRT: severe return visit;
DIA: data-independent acquisition.

## Availability of data and materials

The mass spectrometry proteomics data have been deposited to the ProteomeXchange Consortium (http://proteomecentral.proteomexchange.org) via the iProX partner repository with the dataset identifier.

## Author contributions

Conceptualization: Y.G, C.S, W.S, J.Z, Methodology: J.S, J.W, H.Y, H.S, X.L, Investigation: J.S, Y.Z, H.Y, Visualization: J.W, Supervision: Y.G, C.S, W.S, Writing—original draft: J.W, Y.G, C.S, W.S, Writing—review & editing: J.W, Y.G, C.S, W.S.

## Declarations

### Consent for publication

Not applicable

