## Supplemental Figures for "Urine proteomic characterization of active and recovered COVID-19 patients"

#### Figure S1

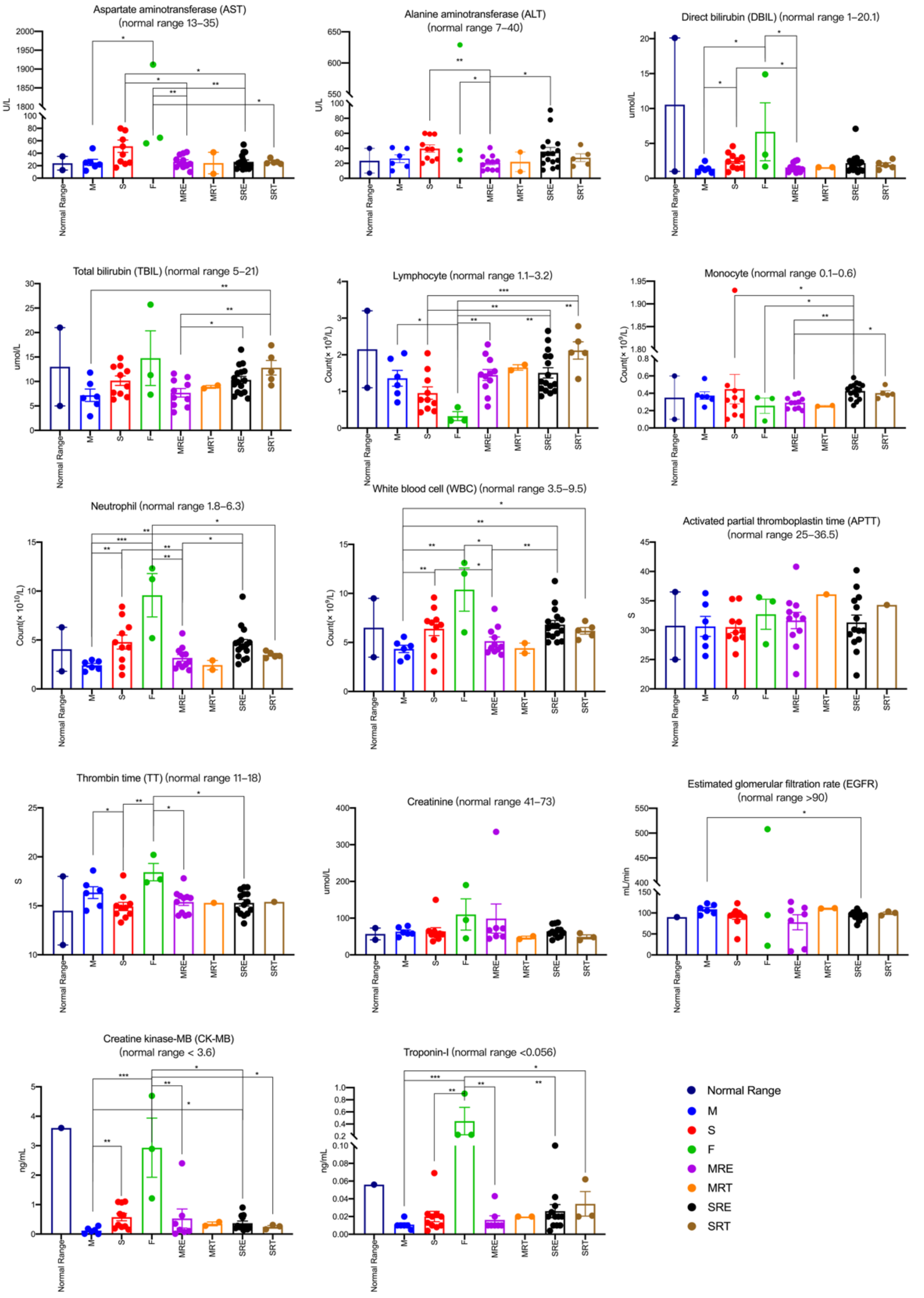


**Figure S1.** The representative 14 clinical parameters in the active and recovery period of COVID-19 patients. The y axis stands for the concentration in serum. The significance indicated by the asterisks. For the clinical parameters, the unpaired two-sided Welch’s t test was performed (p value: *, < 0.05; **, < 0.01; ***, < 0.001).

**Figure S2**

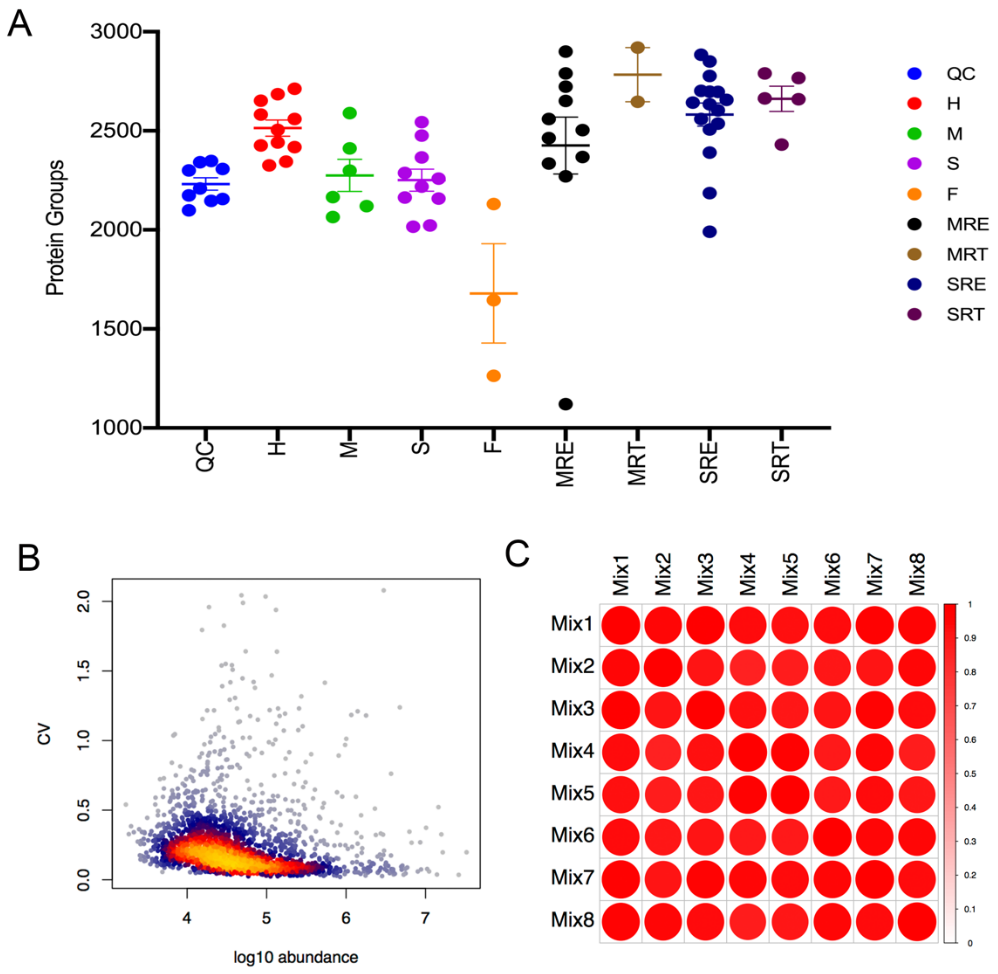


**Figure S2. Protein identification and CV distributions.**

(A). The distribution of numbers of quantified protein groups in the 64 urine samples. Error bars represent multiple independent samples, H (n = 11), M (n = 6), S (n = 10), F (n = 3), MRE (n = 11), MRT (n = 2), SRE (n =16), SRT (n =5).

(B). Coefficient of variation (CV) of the proteomic data is calculated by the proteins quantified in eight quality control (QC) samples using the pooled samples from all urine samples. The X axis denotes the median log10 protein abundance among the QC runs.

(C). The Pearson correlation analysis of the eight QC samples.

**Figure S3**

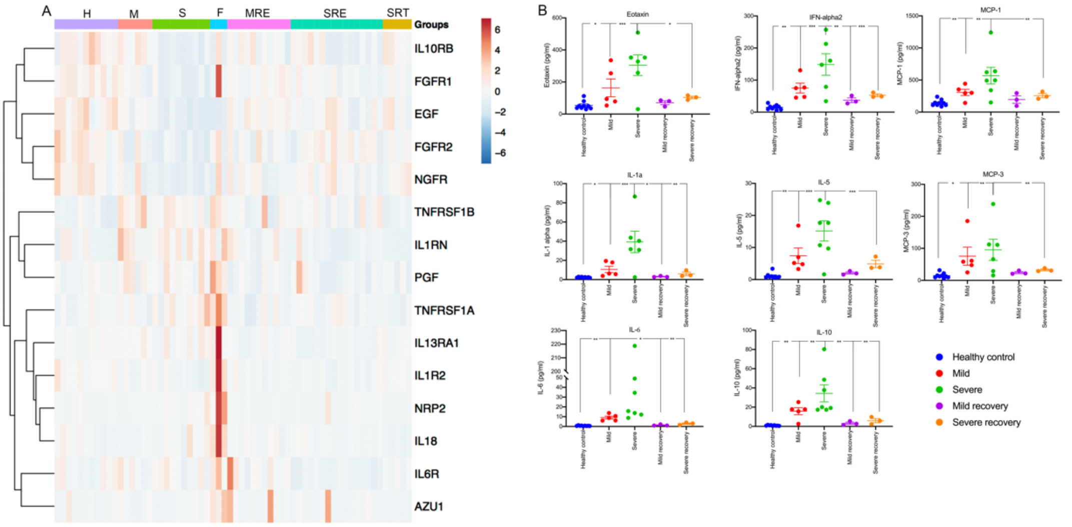


**Figure S3. The associated cytokine proteins.**

(A) Heatmap of 15 differential proteins associated with cytokines identified both in IPA and immport database.

(B). The representative 8 cytokines concentrations in the active and recovery period of COVID-19 patients. The y axis stands for the concentration in serum. The significance indicated by the asterisks. For the clinical parameters, the unpaired two-sided Welch's t test was performed (p value: *, $< 0.05$; **, $< 0.01$; ***, $< 0.001$).

### Figure S4

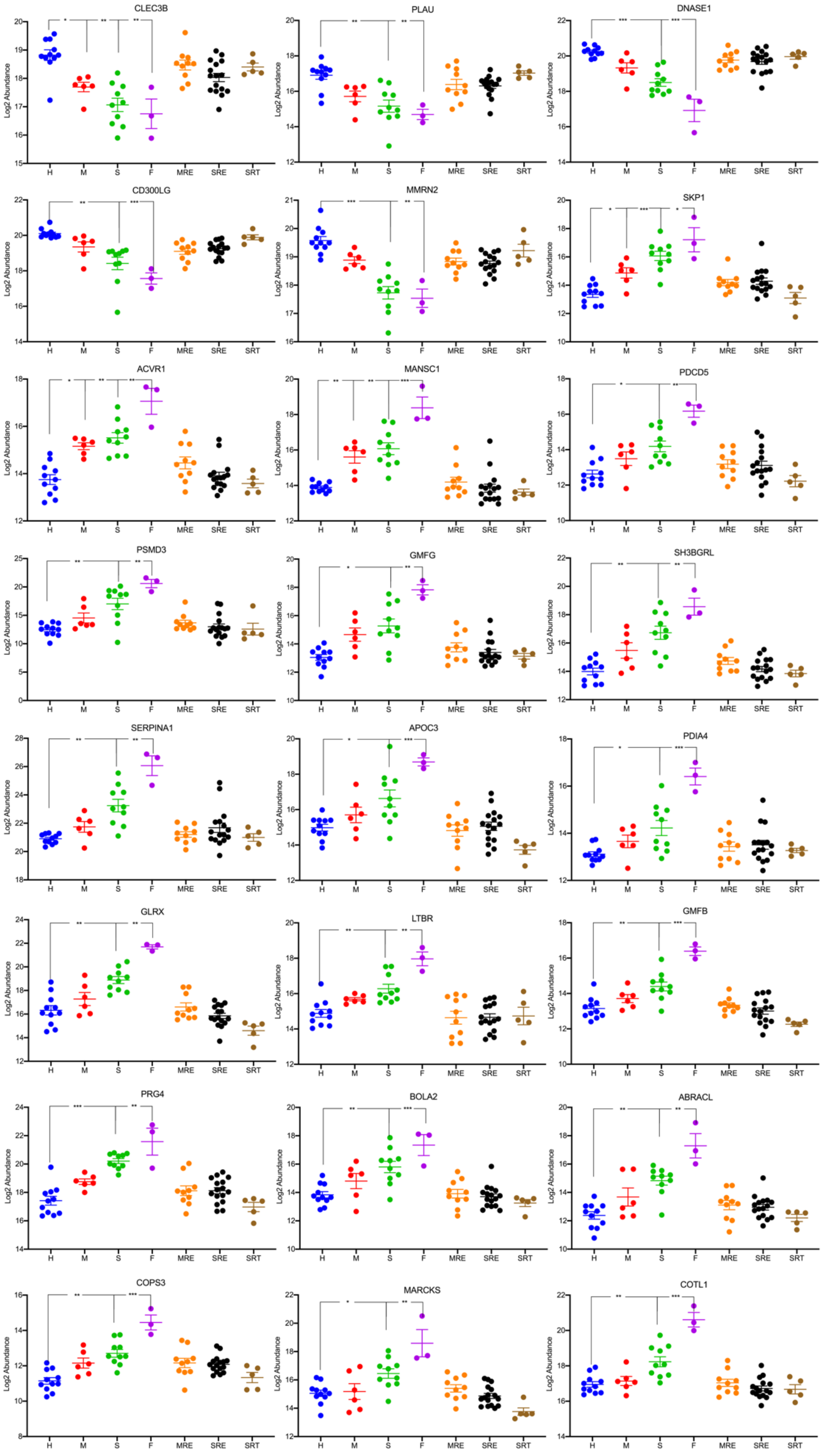
CLEC3B
PLAU
DNASE1
CD300LG
MMRN2
SKP1
ACVR1
MANSC1
PDCD5
PSMD3
GMFG
SH3BGRL
SERPINA1
APOC3
PDIA4
GLRX
LTBR
GMFB
PRG4
BOLA2
ABRACL
COPS3
MARCKS
COTL1
Log2 Abundance
H
M
S
F
MRE
SRE
SRT

**Figure S4. The** 24 differential proteins exhibited great differentiate ability in the three active stages of COVID-19.

**Figure S5**

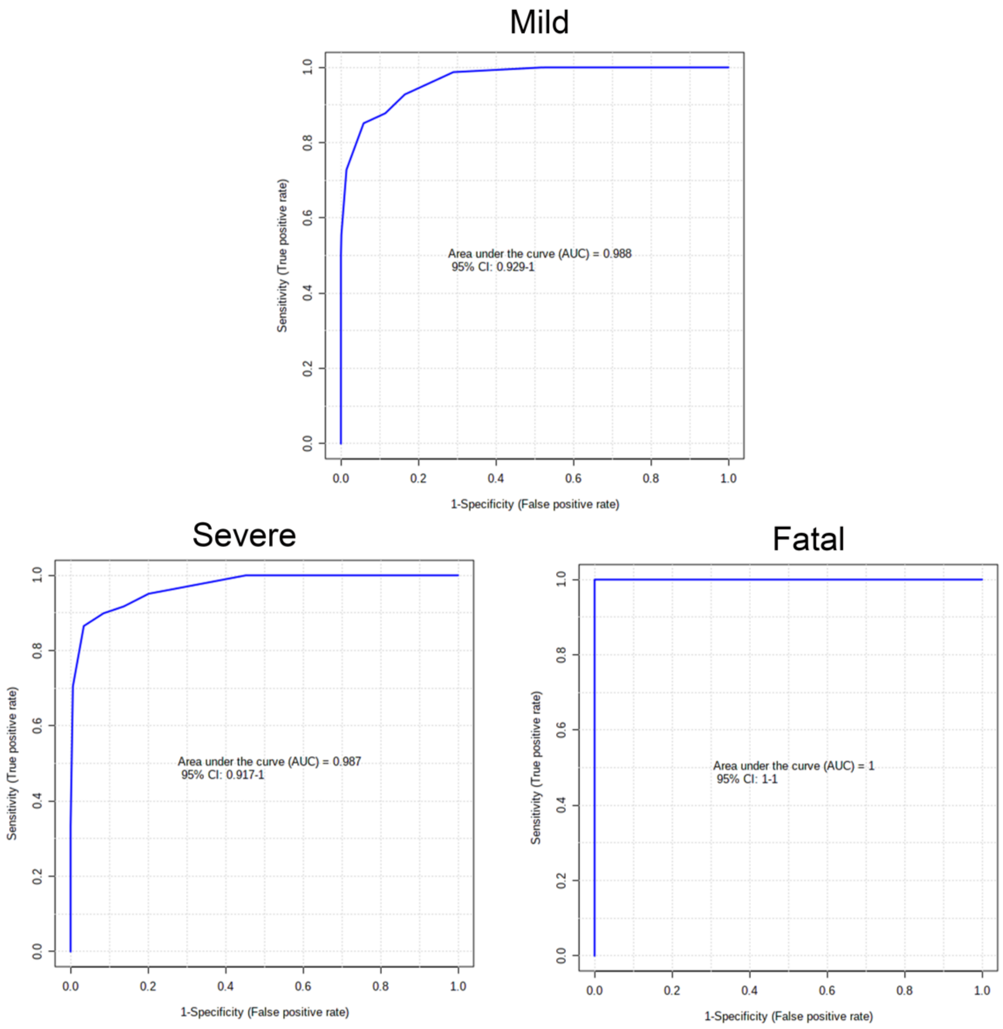


**Figure S5.** Diagnostic performance of KNG1 in the three active stages of COVID-19. The x-axis represents the diagnostic sensitivity of the biomarker panel, and the y-axis represents its diagnostic specificity.
